# Cytokinesis-dependent twisting of HMR-1/Cadherin regulates the first left-right symmetry-breaking event in *Caenorhabditis elegans*

**DOI:** 10.1101/2024.12.12.628066

**Authors:** Mi Jing Khor, Gaganpreet Sangha, Kenji Sugioka

## Abstract

Diverse mechanisms for establishing cellular- and organismal-level left-right (L-R) asymmetry emerged during the evolution of bilateral animals, including cilia-based and actomyosin-dependent mechanisms. In pond snails and *Caenorhabditis elegans*, cell division plays a critical role in regulating both levels of L-R asymmetries. However, the precise mechanism by which cell division breaks cellular-level L-R symmetry remains elusive. Here, we show that cytokinesis-induced cortical flow twists the cell-cell adhesion pattern, which in turn controls the L-R asymmetrical constriction of the contractile ring, thereby breaking the first L-R body symmetry in *C. elegans*. During the second mitosis of *C. elegans* embryos, we discovered the twisting of the HMR-1/cadherin patch at the cell-cell contact site. The HMR-1 patch twisting occurs within a few minutes upon cytokinesis onset, with individual cadherin foci within the patch exhibits directional flow and coalescence. This cell type exhibits chiral cortical flow, characterized by counter-rotational surface flows in the two halves of the dividing cell. We found that this chiral cortical flow plays a critical role in regulating HMR-1 patch twisting by inducing cadherin flow. As the HMR-1 patch twists, the contractile ring preferentially associates with HMR-1 on the right side of the embryo. We demonstrate that HMR-1 patch twisting regulates the L-R asymmetric ring closure. This study uncovers an interplay between three fundamental cellular processes—cell-cell adhesion, cytokinesis, and cell polarity— mediated by cadherin flow, shedding light on cadherin flow’s role in cellular patterning during development.

## Introduction

Bilateral animals typically have a symmetric body plan, but their visceral organs are positioned and shaped asymmetrically along the left-right body axis^1,2^. The mechanisms regulating this animal handedness are hierarchical, with molecular chirality believed to generate cellular chirality, which subsequently establishes organismal-level chirality^3–5^. Prominent examples of cellular-level chirality include the chiral rotation of motile cilia that generates directional fluid flow in the embryos of mice^6,7^, rabbits^6^, frogs^8^, and fish^6^; chiral cell shape changes in fruit flies^9^ and cultured mammalian cells^10,11^; chiral cell migration in chicks^12^; and the chiral orientation of cell division in *C. elegans*^13^ and pond snails^14^. Although key cytoskeletal regulators required for cellular-level chirality have been identified in certain cases, such as Myo1D^15,16^ and Formin^17–19^, the detailed processes and precise mechanisms by which cellular-level L-R asymmetry is achieved remain unclear.

Among the diverse cellular-level chirality described above, the mechanism of cell division-dependent generation is little understood. In snails and *C. elegans*, the direction of cell division along the L-R axis at the 4-8 cell stage and the 4-6 cell stage specifies the chirality of shell coiling and the twisting of the gonad and gut, respectively^13,14^. A common feature in these snails and nematodes is the counter-rotational movement of the cell surface in the two halves of a dividing cell during cytokinesis^20,21^. In *C. elegans*, the actin-nucleating and polymerizing factor Formin is required for this chiral movement of the cell surface^19^. Similar chiral cell surface movements are also observed during cytokinesis in frog zygotes, and drug treatments that disrupt these movements result in abnormal organ chirality^22^. Furthermore, Formin is commonly required for embryonic and/or organismal chirality in snails^17,18^, *C. elegans*^19^, and frogs^17^. Thus, it is tempting to speculate that the cell division-dependent generation of cellular-level chirality is a conserved mechanism invented in a common ancestor of Bilateria. The role of chiral cell surface movement is proposed: it regulates the tilting of the division axis in snails^21^ and *C. elegans*^20^, thereby establishing embryonic chirality. However, the underlying mechanism remains unknown.

In this study, we focused on the first L-R symmetry-breaking event in *C. elegans*, which precedes the above-described division axis tilt along the L-R axis at the 4-6 cell stage. This event occurs during the 2-cell stage cytokinesis, where the contractile ring of the somatic blastomere AB closes toward the right side of the body^23^. Although this cell type also exhibits chiral cell surface movement and dorsal-ventral oriented division axis tilt^24,25^, the rightward contractile ring closure cannot be geometrically explained by a simple tilt in the division axis, as one would expect to see a symmetric contractile ring closure even after the division axis tilt. Clues can be found in previous studies on asymmetric contractile ring closure. Asymmetric contractile ring closure is widely observed in animal zygotes and epithelial tissues, playing a crucial role in regulating cellular patterning^26,27^. In a multicellular context, contractile ring constriction is locally inhibited by adhesion with neighbouring cells, leading to asymmetric closure^28,29^. Therefore, we analyzed the dynamics of the adhesion molecule cadherin during cytokinesis. The *C. elegans* genome encodes a sole classical cadherin HMR-1, which is homologous to vertebrate classical cadherins such as E-cadherin and N-cadherin^30^. By conducting high-resolution 4D live-imaging of cortical flow at the cell surface, HMR-1/cadherin at cell-cell contact, and the contractile ring closure, we identified a potential mechanism for the first L-R symmetry-breaking event in this widely used model organism.

## Results

### Cadherin patch exhibits chiral twisting during L-R symmetry-breaking of cytokinesis

In the *C. elegans* two-cell stage embryo, the anterior AB cell, a somatic founder cell, divides prior to the posterior P_1_ cell (Figure 1A and 1B). During cytokinesis of the AB cell, the contractile ring closes toward the cell-cell boundary with P_1_ and also toward the right side of the body (Figure 1B, schematic and Figure S1A)^23^. Our previous study showed that the asymmetric contractile ring closure requires the neighbouring P_1_ cell: when separated from P_1_, the AB cell exhibited symmetric contractile ring closure (Figure S1B)^31^. We also reported that the attachment of adhesive beads restored the asymmetric closure toward the cell-bead boundary (Figure S1C)^31^. However, after re-analyzing the dataset, we discovered that the L-R asymmetry was not rescued by the bead attachment (Figure S1C-E). These beads were covalently bound to positively charged Rhodamine and adhere to the negatively charged plasma membrane through electrostatic interactions^32^. We reasoned that the L-R asymmetry in contractile ring closure might be generated only through a biological adhesion mechanism involving adhesion molecules, such as cadherins.

**Figure 1.**
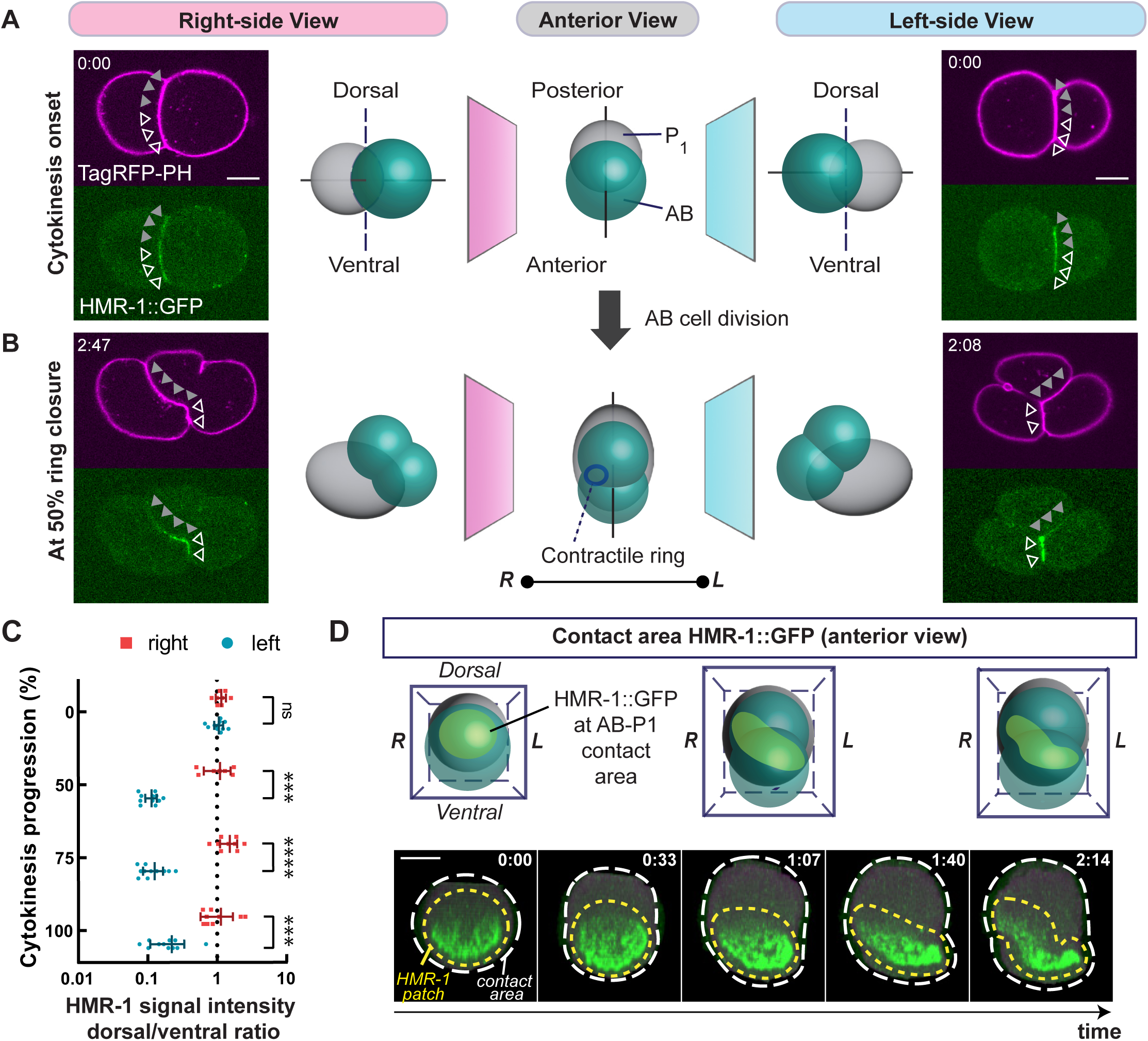
HMR-1/Cadherin exhibits chiral twisting during L-R symmetry-breaking of cytokinesis. (**A**-**B**) Distribution of HMR-1/cadherin at the cell-cell boundary at cytokinesis onset (A) and at 50% contractile ring closure (B). Schematics illustrate 2-to-4 cell stage embryos from the anterior view and lateral views (left and right sides). Micrographs show embryos expressing TagRFP::PH (magenta, plasma membrane) and HMR-1::GFP (green, cadherin). Dorsal and ventral cell-cell boundaries are indicated by gray and white arrowheads, respectively. Upon cleavage furrow formation, the furrow position is defined as the dorsal-ventral boundary. Scale bars: 10 μm. (**C**) Mean dorsal-to-ventral ratio of HMR-1 signal intensity measured on the right and left sides of the embryos. Due to varia-tions in cytokinesis timing, timeseries data were normalized based on the progression of cytokinesis (100%: comple-tion of the contractile ring closure). (**D**) HMR-1 localization at the contact area viewed from the anterior side of the embryo. Micrographs show HMR-1::GFP signals at cell-cell contact areas during AB cell division. Yellow dotted lines and white dotted lines indicate the HMR-1 patch and cell-cell contact area, respectively. Times are relative to cytoki-nesis onset. Scale bar: 5 μm. Error bars: 95% confidence intervals. P-values: Mann-Whitney test (adjusted by the Holm-Sidak method). “****”, “***”, and “ns” indicate p < 0.0001, p < 0.001, and p > 0.05, respectively.

We first characterized the distribution of endogenously tagged HMR-1/cadherin::GFP at the cell-cell contact area during AB cell division. At cytokinesis onset, HMR-1::GFP is uniformly localized at the cell-cell boundary, exhibiting a L-R symmetric distribution pattern (Figure 1A). Indeed, HMR-1::GFP signal intensities at the dorsal (Figure 1A; filled triangles) and ventral halves of the cell-cell boundary (Figure 1A; open triangles) are nearly identical, as evidenced by the dorsal-to-ventral intensity ratio (Figure 1A and 1C). However, as cytokinesis progresses, HMR-1::GFP gradually becomes asymmetric along the dorsal-ventral axis on the left side of the embryo, with ventral signal intensifying compared to the dorsal side (Figure 1B; right figures and Figure 1C). Conversely, the HMR-1 distribution on the right side of the embryos remains relatively uniform along the dorsal-ventral axis (Figure 1B; left figures and Figure 1C). To further investigate the HMR-1 redistribution process, we visualized HMR-1::GFP signals across the entire cell-cell contact area, rather than focusing solely on the left or right surfaces, after computationally removing cytoplasmic and non-junctional HMR-1 signals (Figure 1D, white dotted line; Figure S2A). We defined the region of the cell-cell contact area with higher HMR-1::GFP signal intensity than the surrounding area as “HMR-1 patch” (Figure 1D, yellow dotted line). At cytokinesis onset, the HMR-1 patch is circular, as expected from the geometry of the interface between the two spherical blastomeres. As cytokinesis progresses, we observed that the HMR-1 patch undergoes a counter-clockwise twisting deformation when viewed from the anterior side (Figure 1D, Figure S2B, and Movie S1). The twisting pattern is observed regardless of the orientation of the embryos during imaging, eliminating the possibility of imaging artifacts (Figure S2C). As a result of twisting, HMR-1 molecules are vacated mostly from the dorsal-left side and, to a lesser extent, from the ventral-right side, generating the stronger dorsal-ventral asymmetry on the left side of the body (Figure 1C).

### Chiral cortical flow drives HMR-1 L-R asymmetrical localization

The AB cell is known to exhibit chiral movement of the cell cortex, termed chiral cortical flow, upon cytokinesis onset (Figure 2A and 2B; control)^24,25^. We hypothesized that chiral cortical flow regulates the counter-clockwise deformation of the cadherin patch. To test this hypothesis, we experimentally reduced cortical flow chirality. Previous studies have shown that the RhoA signaling regulates chiral cortical flow^20^. Thus, we performed feeding RNAi knockdown of ECT-2/RhoGEF, an RhoA activator. In order to obtain varying effects on chiral cortical flow, we have conducted RNAi feeding for different durations: 4, 6, and 8 hours. These treatments are all mild knockdowns of ECT-2, with cytokinesis completing in all samples. With longer RNAi feeding durations, we observed a proportional reduction in chiral cortical flow strength (Figures 2B-C and Movie S2). Next, we compared the HMR-1 patch twisting pattern at the cell-cell contact area between control embryos and embryos subjected to 4, 6, and 8 hours of *ect-2(RNAi)*. We found that the level of HMR-1 twisting, measured by the angle of the line connecting the dorsal and ventral centroids of the HMR-1 patch (Figure S3A-B), gradually decreased as the RNAi duration increased (Figure 2D-E and Movie S1). Consistently, the dorsal-ventral asymmetry in the HMR-1 signal was also lost on the left surface of the embryo (Figure S3C). These results suggest that chiral cortical flow is required for HMR-1 patch twisting.

**Figure 2.**
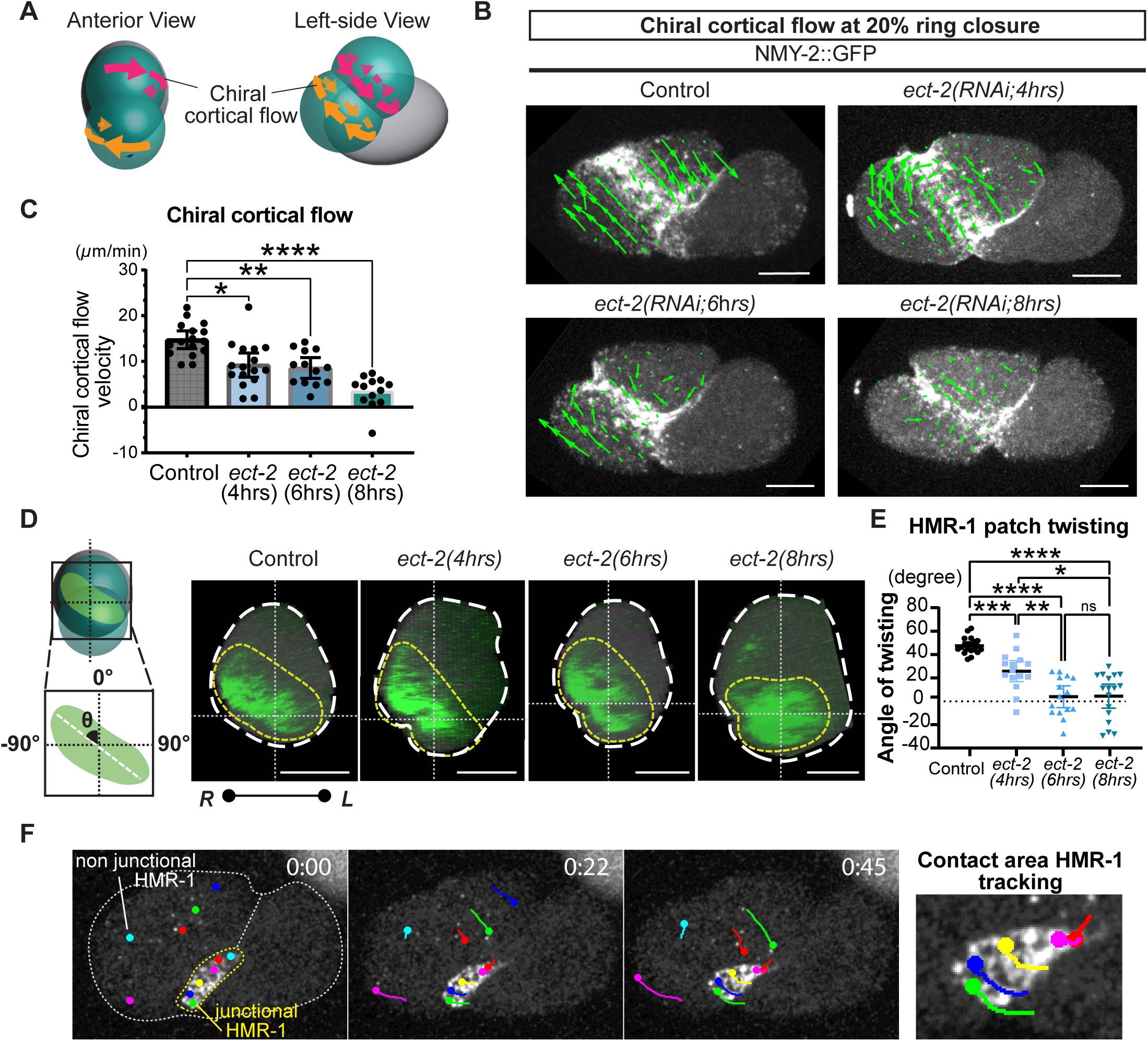
Chiral cortical flow regulates twisting of the HMR-1 patch at the cell-cell contact. (**A**) The AB cell exhibits chiral cortical flow during cytokinesis, where the two halves of the dividing cell counter-rotate relative to each other. (**B**) Cortical flow during cytokinesis imaged using non-muscle myosin II::GFP (NMY-2) in control and *ect-2(RNAi)* embryos. Cortical flow vectors (green arrows) were estimated by particle image velocimetry (PIV). Scale bar: 10 µm (**C**) Mean chiral cortical flow measured over 4 time points (22 seconds) upon reaching 20% ring closure in control and *ect-2(RNAi)* embryos. (**D**-**E**) HMR-1 patch at the cell-cell contact area in control and *ect-2(RNAi)* embryos at 50% contractile ring closure (D). Yellow dotted lines indicate the outlines of the HMR-1 patch. White dotted lines denote the D-V and L-R body axes, with horizontal line also marking the dorsal-ventral boundary (defined by the cleavage furrow position). Scale bar: 10 µm. Mean HMR-1 patch orientation relative to the D-V axis is shown in the graph (E; See Methods for detail). (**F**) Tracking of HMR-1::GFP during cytokinesis. Trajecto-ries of two populations of HMR-1, at the cell surface and cell-cell contact, are shown. Contact area HMR-1 foci are visible as the embryo is viewed from the ventral left side. Times are relative to the onset of HMR-1 patch deforma-tion. Error bars: 95% confidence intervals. P-values: Brown-Forsythe and Welch ANOVA, followed by the Dunnett T3 multiple comparison test. “****”, “***”, “**”, “*”, and “ns” indicate p < 0.0001, p < 0.001, p < 0.01, p < 0.05, and p > 0.05, respectively.

How might chiral cortical flow regulate the twisting of the HMR-1 patch? We previously showed that extracellular particles coating the plasma membrane of the AB cell exhibited chiral flow along with the cortical myosin^24^. This observation suggests that transmembrane proteins move together with the actomyosin cortex. Consistently, non-junctional HMR-1 in zygotes exhibits cortical flow^33^. As expected, during cytokinesis of the AB cell, we observed that non-junctional HMR-1 foci at the cell surface exhibited chiral cortical flow (Figure 2F; non-junctional HMR-1, Movie S3). Furthermore, we observed the corresponding chiral movement of HMR-1 foci at the cell-cell contact area (Figure 2F; junctional HMR-1, Movie S3). This observation, along with the results of *ect-2(RNAi)*, suggests that chiral cortical flow regulates the asymmetric movements of HMR-1. The redistribution of cadherin on the plasma membrane can be generally explained by either endocytic recycling^34^, membrane diffusion^35^, or directed transport known as cadherin flow^37^. Our observation suggests that cadherin flow underlies this phenomenon. Consistent with this idea, multiple myosin foci coalesce during their movement, which is accompanied by an increase in the maximum HMR-1 signal intensity at the cell-cell contact area—an effect that cannot be explained by other mechanisms of cadherin redistribution (Figure S4A-B). Taken together, these results suggest that chiral cortical flow directs the L-R asymmetric flow of HMR-1 foci on the membrane, generating the twisted pattern of the HMR-1 patch.

### HMR-1 patch twisting is required for L-R asymmetric contractile ring closure

In the in vitro adhesive bead experiments shown in Figure S1, where adhesive molecules were covalently bound to the bead surface and thus rendered immobile, L-R asymmetric contractile ring closure was not observed. In contrast, the directed membrane movement of HMR-1 foci and the formation of their L-R asymmetric pattern may account for the L-R asymmetric contractile ring closure in vivo. To investigate this, we analyzed the dynamics of contractile ring in the absence of HMR-1 patch twisting. In control embryos, the rightward shift of the contractile ring gradually begins and reaches to its peak at approximately 70% ring closure (Figure 3A-C and Movie S4). However, this rightward shift was progressively lost with increasing duration of *ect-2(RNAi)* treatment (Figure 3A-C and Movie S4). In some *ect-2(RNAi)* embryos, we observed the loss or reversal of the HMR-1 patch twisting (Figure 2E). Strikingly, we also observed the corresponding loss or reversal of the contractile closure asymmetry in these embryos (Figure 3D). By using data obtained from control and *ect-2(RNAi)* embryos, we found a positive correlation between the degree of HMR-1 patch twisting and the rightward shift of the contractile ring (Figure 3E). These results suggest that HMR-1 patch twisting regulates L-R asymmetric contractile ring closure.

**Figure 3.**
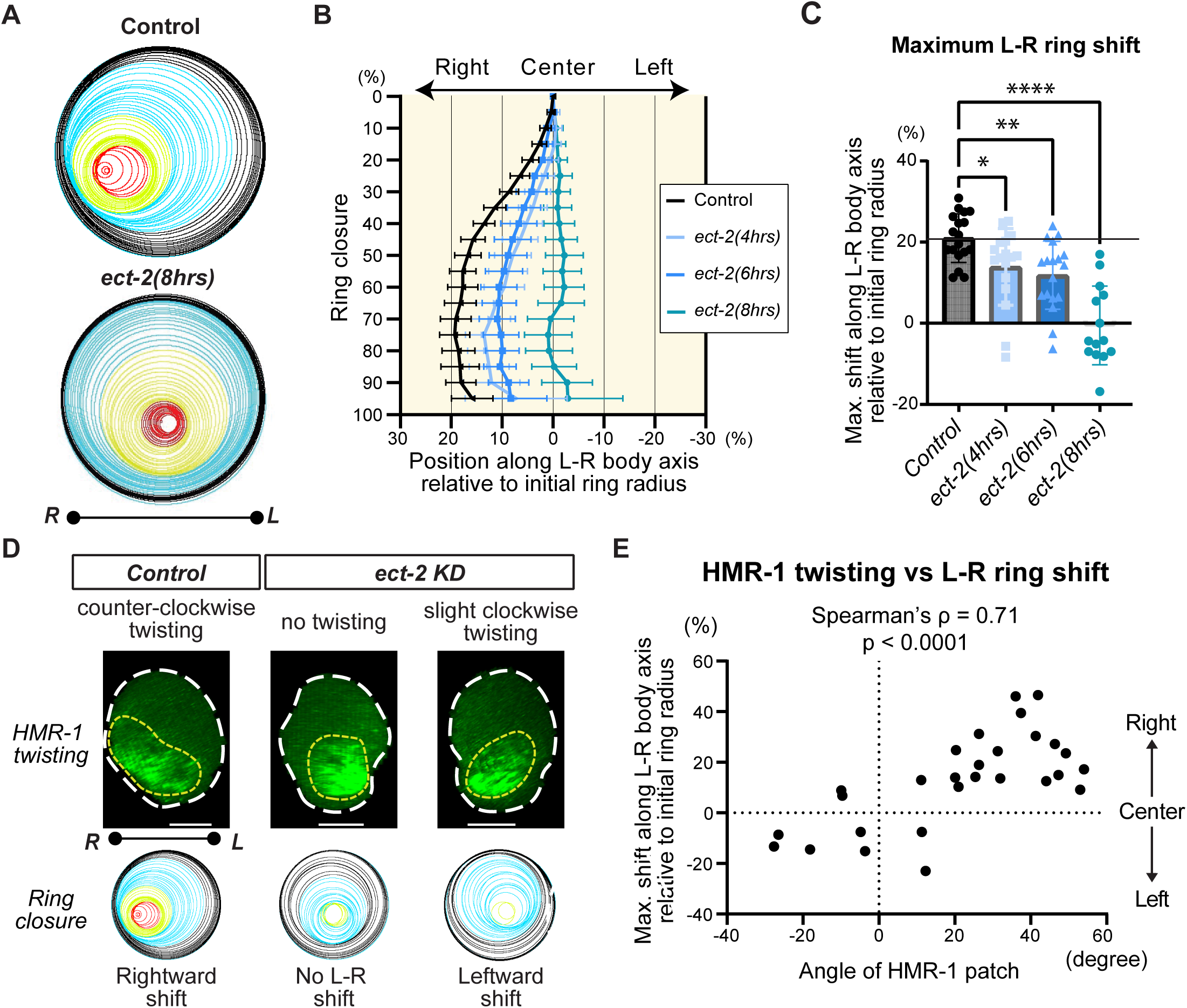
HMR-1 patch twisting is associated with the L-R contractile ring shift. (**A**) Contractile ring trajectory of control and *ect-2* knockdown embryos. The color code indicates the time after cytokinesis onset. (**B**) The shift in the position of the contractile ring along the L-R axis during cytokinesis. The x-axis represents the deviation of the centroid of the contractile ring as a percentage relative to the ring radius at cytokinesis onset. (**C**) Maximum shift of the contractile ring along the L-R axis. Error bars: 95% confidence inter-vals. P-values: Brown-Forsythe and Welch ANOVA, followed by the Dunnett T3 multiple comparison test. “****”, “**”, “*” indicate p < 0.0001, p < 0.01, p < 0.05, respectively. (**D**) HMR-1 twisting and ring closure patterns in individual samples. Notably, in some *ect-2* KD embryos, the orientation of the HMR-1 patch twisting and ring closure were reversed. (**E**) Relationship between the level of HMR-1 twisting (represented by the angle of the HMR-1 patch) and the maximum L-R ring shift. Error bars indicate 95% confidence intervals.

Now that chiral cortical flow, HMR-1 patch twisting, and L-R asymmetric contractile ring closure are all correlated, we sought to understand the hierarchy among these relationships. To investigate this, we conducted *hmr-1(RNAi)*. As previously reported, HMR-1 is not the sole adhesion molecule mediating early embryonic cell-cell adhesion in *C. elegans*, but works with SAX-7/L1CAM and other unidentified adhesion molecules^36^. About 14% *hmr-1(RNAi); sax-7(eq1)* embryos exhibited abnormal AB cell arrangement in which the contractile ring was displaced from the AB-P_1_ boundary (4 out of 29). The subsequent analyses were conducted using samples that still maintain the normal AB and P_1_ cell arrangement. We found that the contractile ring exhibited a reduced rightward shift in these *hmr-1(RNAi); sax-7(eq1)* embryos (Figure 4A-B and Movie S4). These embryos retained the normal levels of chiral cortical flow (Figure 4C). Furthermore, the positive correlation between chiral cortical flow and L-R asymmetric contractile ring closure, which was observed in the control, was lost in *hmr-1(RNAi);sax-7(eq1)* (Figure 4D). These results suggest that while chiral cortical flow is required, it is not sufficient to induce L-R asymmetric contractile ring closure. Rather, it requires downstream adhesion molecule, such as HMR-1 and SAX-7, to mediate the rightward shift of the contractile ring.

**Figure 4.**
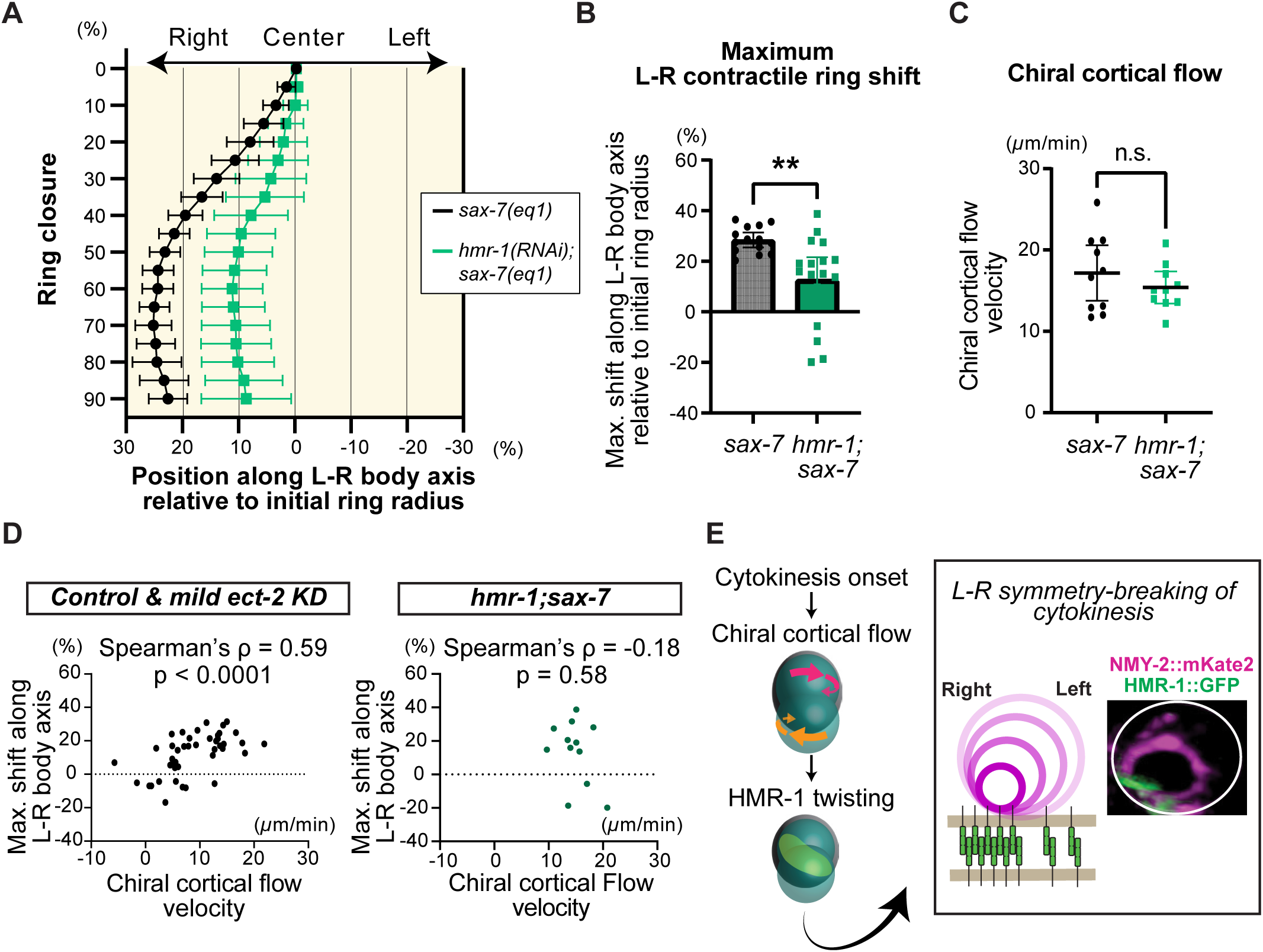
HMR-1 is required for L-R symmetry-breaking during cytokinesis. (**A**) The shift in the position of the contractile ring along the L-R axis during cytokinesis in *sax-7(eq1)* and *hmr-1(RNAi)*;*sax-7(eq1)* embryos. *sax-7(eq1)* homozygous mutants are viable, whereas *hmr-1(R-NAi);sax-7(eq1)* embryos exhibit100% embryonic lethal and fail gastrulation. See Figure 3B for comparison with wild-type control. (**B**) Maximum shift of the contractile ring along the L-R axis. (**C**) Chiral cortical flow is not altered by *hmr-1(RNAi)*. (**D**) Correlation between chiral cortical flow velocity and the maximum L-R shift of the contractile ring. (**E**) A proposed model of the first L-R symmetry-breaking in *C. elegans*. Cytokinesis induces chiral cortical flow in a formin-dependent manner, which causes twisting of the HMR-1 patch. The L-R asymmet-ric HMR-1 then preferentially interacts with the contractile ring on the right side of the embryo (as shown in the micrograph). This local adhesion inhibits contractile ring closure, resulting in a rightward shift of the contractile ring. Error bars indicate 95% confidence intervals. Error bars: 95% confidence intervals. P-values: Welch’s t-test. “**” and “ns” indicate p < 0.01 and p > 0.05, respectively.

## Discussion

In this study, we documented the mechanism underlying the left-right asymmetric closure of the contractile ring, the first L-R symmetry-breaking event in *C. elegans*. Although synthetic adhesives, which mediate adhesion through electrostatic interactions, can induce asymmetric contractile ring closure toward the adhesive surface, we found that they were unable to induce the L-R asymmetry observed in vivo. Through investigation of the in vivo adhesion pattern, we discovered that HMR-1/cadherin foci undergo L-R asymmetric flow, leading to the twisted pattern of their clusters. We demonstrated that chiral cortical flow regulates HMR-1 patch twisting. Furthermore, we showed that HMR-1 patch twisting is associated with L-R asymmetric contractile ring closure and that a reversal of L-R asymmetry can be induced. Finally, we showed that *hmr-1(RNAi)* reduced L-R asymmetric contractile ring closure without altering the chiral cortical flow.

Based on the obtained results, we propose a mechanism for the L-R symmetry-breaking of cytokinesis (Figure 4E). First, cytokinesis induces chiral cortical flow in a RhoA signaling-dependent manner. Chiral cortical flow then transports HMR-1 foci asymmetrically along the cell-cell contact membrane, leading to the generation of cadherin flow and twisting of HMR-1 patch. The contractile ring preferentially associates with HMR-1 on the right side of the embryo (Figure 4E; micrograph). As demonstrated in our previous study, adhesion can locally inhibit ring closure in the AB cell by limiting the cortical flow directed toward the contractile ring (which is orthogonal to the chiral cortical flow)^31^. Consequently, the contractile ring closure is more restricted on the right side of the embryo, resulting in a rightward shift of the contractile ring.

Cortical flow is known to directionally transport cadherin in mammalian and zebrafish cultured cells during interphase and has a critical role in remodeling adhesion complex^37–39^. However, the role of this cadherin flow during cell division remained unclear. The evidence we present is expected to open new lines of inquiry into how cytokinesis, adhesion, and cell polarity—three fundamentally important cellular processes—interplay during development via cadherin flow. A previously known example of interplay between these cellular processes is the cytokinesis of *Drosophila* epithelial cells. During epithelial division, adherens junctions at the apical side physically restrict contractile ring constriction, inducing basal-to-apical contractile ring closure, which is critical for maintaining epithelial polarity^28,40^. Due to this physical restriction, the completion of cytokinesis requires the disengagement of adherens junction from the contractile ring^28^. This adhesion disengagement is coupled with the reduction of E-cadherin level at the cleavage furrow, which induces cortical flow in the neighbouring cell^41^. Thus, in this context, cortical flow is downstream of cadherin and does not induce cadherin flow^41^. During cytokinesis of *C. elegans* zygotes, non-junctional HMR-1/cadherin foci exhibit flow along with myosin^33^, but the role of this cadherin flow remained unclear. Our study suggests that cortical flow during cytokinesis induces cadherin flow that in turn generates cellular-level chirality. We believe that synthetic adhesives used in Figure S1 could not induce L-R asymmetric ring closure because they are rendered immobile and, unlike cadherin flow, cannot translate chiral flow into an L-R asymmetric adhesion pattern. Overall, this study underscores the significance of the cadherin flow in regulating cell division-dependent chiral morphogenesis.

Our analysis of the cadherin redistribution process is limited in spatial resolution, leaving the molecular changes to HMR-1/cadherin largely unclear. A circular cluster of HMR-1::GFP foci, HMR-1 patch, forms at the cell-cell contact immediately after the cytokinesis of the previous cell division around 13 min prior to the cytokinesis of the AB cell. Since trans-dimers of cadherin can form immediately after cell-cell contact^42^, and the separated AB and P_1_ cells can re-establish adhesion almost instantly after contact^32^, we speculate that the HMR-1 patch consists of trans-dimers of HMR-1 molecules. We observed rapid HMR-1 foci movement and an increase in HMR-1::GFP signal intensity along the direction of twisting. This observation suggests that HMR-1 trans-dimers move along the plasma membrane. The directional nature of this movement rules out the possibility of random cadherin diffusion^34^ mediating this process. The contribution of endocytosis, another regulatory mechanism of cadherin redistribution^34^, is also deemed unlikely, as we directly observed the movement of HMR-1 foci (Movie S1) and their displacement from one place to another takes on the scale of seconds (Figure S4). In contrast, endocytosis-dependent turnover typically occurs on the scale of minutes^34,43^. It is of great interest to explore whether cortical flow-dependent redistribution of cadherin can alter the molecular activity of cadherin, such as cis-clustering^44^, in a mechanosensitive manner.

Among the diverse mechanisms regulating organismal-level L-R asymmetry in bilaterians^5^, motile cilia-dependent activation of L-R asymmetric Nodal signaling has garnered significant attention due to its conservation in mammals^7^ and its association with ciliopathies and heterotaxy syndromes^45,47^. *C. elegans* lost Nodal signaling during evolution, however, the cell division-dependent mechanism and Nodal signaling are not mutually exclusive. Snails rely on cell division for chiral morphogenesis but somehow activate L-R asymmetric Nodal signaling^49^. Frogs are known to rely on motile-cilia dependent L-R asymmetric Nodal signaling^8^, but perturbation of zygote chiral cell surface flow during cytokinesis results in organ chirality failure^22^. While independent co-option might be a possibility, these three organisms all require formin gene for embryonic and/or organismal chiral morphogenesis^17–19^. Thus, cell division-dependent chiral morphogenesis might be an ancient program deeply encoded across bilaterians. While the significance of the 2-cell stage L-R asymmetric contractile ring closure in later development has not been determined—unlike the L-R asymmetric division axis tilt at the 4-6 cell stage^13,23^—our study demonstrates how adhesion translates chiral cortical flow into L-R asymmetric contractile ring closure. Since chiral cell surface movement is observed in snails and frogs, a similar interplay between cortical flow and cadherin may play a role in cell division-dependent chiral morphogenesis in other organisms.

## Acknowledgements

We thank *Caenorhabditis* Genetics Center (funded by the NIH Office of Research Infrastructure Programs; P40 OD010440) for sharing worm strains and the Sugioka lab members for general discussions. This work was supported by the Canadian Institutes of Health Research (Project Grant; PJT-169145), Government of Canada’s New Frontiers in Research Fund (NFRFE-2019-00310), and the Michael Smith Health Research BC (Scholar Award; SCH-2020-0406) to K.S.

## Author contributions

**M.K.**: Formal analysis, Investigation, Writing-Original Draft, Validation, Visualization. **G.S.**: Formal analysis, Investigation, Validation, Visualization. **K.S.**: Conceptualization, Formal analysis, Investigation, Methodology, Validation, Writing-Original Draft, Writing-Review & Editing, Supervision, Funding acquisition.

## Supplementary Movie legends

Movie S1. HMR-1::GFP localization at the cell-cell contact in control and *ect-2(RNAi, 8hrs)* embryos. The image is en face view of the cell-cell contact and a view from the dorsal anterior side.

Movie S2. Chiral cortical flow in the control and *ect-2(RNAi, 8hrs)* AB cell. Five time points of live imaging used for quantification are shown.

Movie S3. Chiral cadherin flow in control AB cell. Individual HMR-1 foci were tracked by different colors.

Movie S4. L-R asymmetric contractile ring closure in control, *ect-2(RNAi, 8hrs)*, and *hmr-1(RNAi);sax-7(eq1)*.

## STAR Methods

### Contact for Reagent and Resource Sharing

Further information and requests for resources and reagents should be directed to and will be fulfilled by the Lead Contact, Kenji Sugioka.

### Experimental Model and Subject Details

All *C. elegans* strains were cultured at 22.5°C using the standard method^46^. The following transgenes were used: *cp13*[*nmy-2*::GFP + LoxP] (non-muscle myosin II; knock-in)^48^, *cp69*[*nmy-2*::mKate2 + LoxP] (knock-in)^50^, *cpIs56*[*mex-5*p::TagRFP-T::PLC(delta)-PH::*tbb-2* 3’UTR + *unc-119*(+)] (plasma membrane marker)^51^, *or1940*[GFP::*sas-7*] (centriole marker; knock-in)^52^, *ltIs37*[*pie-1*p::mCherry::*his-58* + *unc-119*(+)] (chromosome marker)^53^, *cp21*[*hmr-1*::GFP + LoxP] (HMR-1 cadherin, knock-in)^54^. RNAi knockdown of HMR-1 was conducted in *sax-7*(*eq1*), a genetically null mutant of *sax-7*/L1CAM^55^.

### Method Details

#### RNAi

Feeding RNAi was performed at 22.5°C using the standard method^56^. For *ect-2* knockdown, L4 larvae or young adults were cultured on freshly prepared feeding RNAi plates and imaged at 4, 6, and 8 hours. For *hmr-1* knockdown, L1 starved larvae were prepared by bleaching gravid adults, followed by culturing embryos in sterile M9 buffer for over 24 hours. The synchronized L1 larvae were then transferred to freshly prepared feeding RNAi plates (day 1) and imaged on day 3.

#### Live Imaging

Embryos were obtained by dissecting gravid adults in a droplet of 10-12 *µ*l of refractive index-matching medium^57^, consisting of 30% iodixanol diluted in egg salt buffer and 30 *µ*m diameter plain polystyrene beads, on a coverslip as described before^31^. After gently mounting the coverslip onto a slide glass, three edges of the coverslip were sealed with petroleum jelly (Vaseline), leaving one edge open.

The prepared samples were imaged using an Olympus IX83 microscope (Olympus/Evident), equipped with a spinning-disk confocal unit CSU-W1 (Yokogawa), a scientific CMOS camera Prime95B (Photometrics), a piezoelectric stage NANO-Z (Mad City Labs), a silicon immersion objective UPLSAPO60XS2 (NA1.3, 60X; Olympus/Evident), and a beam splitter Optosplit II (Cairn Research), all controlled by Cellsense Dimension software (Olympus/Evident). Silicone immersion oil (Z81114; refractive index: 1.406 at 23 °C; Olympus/Evident) was used as the immersion medium. Simultaneous two-color imaging was performed using 488 nm and 561 nm diode-pumped lasers, with settings of 150 ms exposure time, 1 µm Z-step size, 31 slices per frame, and a 5.57 second frame interval. Sample collection typically began during metaphase. The obtained 4D imaging data were deconvoluted using a constrained iterative and advanced maximum likelihood algorithm (iteration: 5), using Cellsens Dimension software.

#### Quantification of contractile ring dynamics

We measured contractile ring closure dynamics using our previously described method^55^. When we define the radius of the contractile ring at cytokinesis onset and at a certain time point as *R_0_* and *R_t_*, respectively, the degree of ring closure at time point *t*, *R_closure_(t)*, is defined as follows:

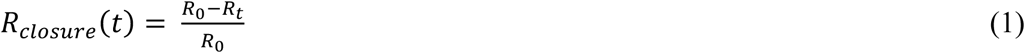

Cytokinesis is completed when *R_closure_(t)* reaches 1. We used this value as an indicator of cytokinesis progression to align the time series data.

To measure the shift of the contractile ring from its initial center, the distance between the centroids of the initial contractile ring and the contractile ring at time *t* was measured. In most of the study, we measured the distance only along the L-R axis. In Figure S1, we also measured the distance along the apico-basal axis. The distance was defined as Qt, and the shift of the contractile ring along the L-R axis or apico-basal axis was defined as follows:

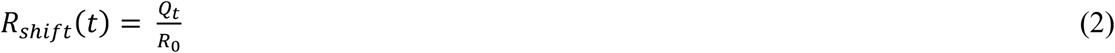

After 4D-imaging of a worm strain expressing non-muscle myosin II::GFP, a centriole marker, a chromosome marker, and a plasma membrane marker, the obtained images were deconvoluted as described above. The deconvoluted images were then further processed using Gaussian blur (sigma = 2) and an image J^58^ plug-in, “attenuation correction” (opening = 3, reference = 15)^59^, to facilitate later image segmentation. For these images, the anterior-posterior axis was determined based on the position of the AB and P_1_ cells. The nuclei of the dorsally located ABp and ventrally located EMS cells were visually marked. Using the Image J plug-in Clear Volume^60^, we adjusted the orientation of volumetric images so that the ABp and EMS nuclei aligned vertically. The en face view of the contractile ring, as seen from the relatively anterior side, was then generated for each time point. The contractile ring regions were computationally detected by image segmentation using an Image J plug-in MorphoLibJ^61^. *R_0_*, *R_t_*, and *Q_t_* were then measured using ellipse fitting of the segmented binary images. The ring trajectory images were generated using an in-house image J macro described before^59^.

For the data in Figure S1, volumetric images were oriented such that the AB cell and the cell or the bead near the contractile ring were aligned vertically. *R_0_*, the contractile ring diameter at a later stage of cytokinesis (60∼80% cytokinesis progression), and the shift of the contractile ring along different axes were measured manually.

#### Quantification of the dorsal-to-ventral ratio of HMR-1 signal intensity

For this measurement, we used laterally oriented embryos expressing HMR-1::GFP and a plasma membrane marker, facing either the left or right side toward the objective lens. Image deconvolution was not performed for this analysis. Before cleavage furrow formation, the dorsal and ventral regions of the AB-P_1_ boundary were simply defined by dividing it into top and bottom halves. After furrow formation, the dorsal and ventral parts of the cell-cell boundary were separated by the furrow position. We measured the average signal intensity of HMR-1::GFP for each dorsal and ventral region using a single z-slice at 0%, 50%, 75%, and 100% cytokinesis progression. The z-slice was selected as the plane closest to the objective lens (the surface of the embryo) where the plasma membrane marker was clearly visible in both the dorsal and ventral regions. The obtained dorsal and ventral cell-cell boundary HMR-1::GFP intensities were defined as *D_avg_* and *V_avg_*, respectively. We also measured the cytoplasmic HMR-1::GFP signal intensity in the AB cell, defined as *cyto_avg_*. The dorsal-to-ventral HMR-1::GFP signal intensity ratio was calculated as follows:

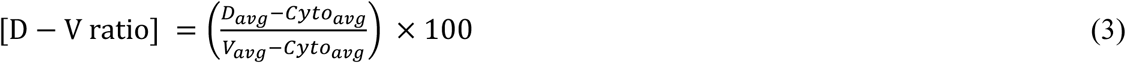

#### Visualization of the HMR-1 patch at the cell-cell contact area

We collected 4D imaging data of a strain expressing HMR-1::GFP and a plasma membrane marker using the acquisition setting described above. All subsequent processing was performed using Image J. To computationally remove HMR-1::GFP signals outside the cell-cell contact area, we first processed the plasma membrane maker signals (Figure S2). For the plasma membrane marker signals, Gaussian blur (sigma = 2) and “attenuation correction” (opening = 3, reference = 15) were applied to facilitate image segmentation. Using the image J plug-in MorphoLibJ, 3D morphological segmentation of AB and P_1_ cells was performed for each timepoint. The segmented binary images for AB and P_1_ cell regions were slightly expanded using dilation function 10 times. These dilated binary images for AB and P_1_ cell regions were then subjected to the Image Calculator (AND operation) to eliminate areas outside the overlapping region (Figure S2). Using these cell-cell contact area binary images, the HMR-1::GFP channel was processed with the Image Calculator (AND operation) to remove HMR-1::GFP signals outside the contact-area. En face views of the contact-area HMR-1::GFP signals were then generated using the ImageJ plug-in Clear Volume. Note that this represents the view from the anterior and slightly dorsal side, with the D-V axis aligned vertically in the image.

#### Quantification of the HMR-1 patch twisting

The HMR-1 patch twisting shown in Figure 2E was measured using the en face view of the contact area HMR-1::GFP signal at 50% cytokinesis progression, as described above (Figure S3). The contact area HMR-1::GFP signal was divided into dorsal and ventral regions based on the position of the cleavage furrow. Segmentation of the dorsal and ventral HMR-1 regions was performed using the thresholding algorithms “IsoData” or “MaxEntropy”, selecting the method that best represented the original images. For each dorsal and ventral region of the segmented area, centroids were determined by applying ellipse fitting. The coordinates of the dorsal and ventral centroids were defined as (x_dorsal_, y_dorsal_) and (x_ventral_, y_ventral_), respectively. Finally, the angle between the line passing through the dorsal and ventral centroids and the vertical D-V axis was calculated as follows to quantify the angle HMR-1 twisting:

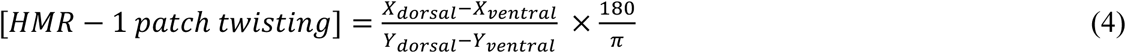

#### Quantification of chiral cortical flow

We collected 4D imaging data of a strain expressing non-muscle myosin II::GFP using the acquisition setting described above. The obtained images were flattened by maximum intensity projection of the 10 surface z-slices closest to the objective lens. Visualization of cortical flow in Figure 2B was performed using PIV lab software^62^. For quantification of chiral cortical flow, the images were rotated to align the cleavage furrow with the vertical axis of the image. Using these images, myosin foci in AB cells were tracked with “Manual Tracking” plug-in of ImageJ. To minimize the effects of AB cell body movement, tracking was performed over five time frames starting from 20% cytokinesis progression. Five foci were tracked for each side of the cortex, separated by the cleavage furrow in the dividing cells. The average velocity of the foci along the vertical axis was calculated to quantify the cortical flow along the axis aligned with the cleavage furrow. We defined the velocities for the left and right sides as v_y, left_ and v_y, right_, respectively. The chiral cortical flow velocity (degree of counter-rotation of flow) was then measured as follows :

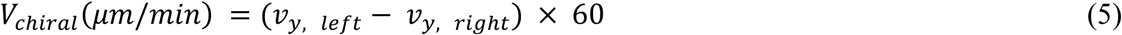

#### Statistics and reproducibility

Statistical analyses were performed using Prism 9 (Graphpad). Welch’s t-test was conducted for comparisons of means between two groups. For comparison of data in Fig. 1C, multiple Mann-Whitney tests were conducted, with P-values adjusted using the Holm-Sidak method. Brown-Forsythe and Welch ANOVA, followed by the Dunnett T3 multiple comparison test, was performed for comparisons of means across multiple groups. Error bars represent the 95% confidence interval. The p-values, p < 0.0001, p < 0.001, p < 0.01, p < 0.05, and p > 0.05 are represented by the symbols, “****”, “***”, “**”, “*”, and “ns”, respectively. Correlational analysis was performed using Spearman’s correlation test. Sample sizes: Fig. 1C (*n* = 11, 8, 11, 7, 11, 9, 11, 9 from top to bottom), Fig. 2C (*n* = 16, 13, 16, 13), Fig 2E (*n* = 20, 15, 15, 17 from the left), Fig. 3B (*n* = control: 20, *ect-2* (4Hrs): 17, *ect-2* (6Hrs): 18, *ect-2* (8Hrs): 13), Fig. 3C (*n* = control: 18, *ect-2* (4Hrs): 17, *ect-2* (6Hrs): 18, *ect-2* (8Hrs): 14), Fig. 3E (*n* = 28), Fig. 4A & B (*n* = control: 14, *hmr-1*(*RNAi*); *sax-7(eq1)*: 17), Fig. 4C (*n* = control: 10, *hmr-1*(*RNAi*); *sax-7(eq1)*: 11), Fig. 4D (n = 43, 12 for left and right panel, respectively), Fig. S1D (*n* = 11, 7 from left), Fig. S1E (*n* = 11, 7), Fig. S2B (*n* = 6), Fig. S4B (*n* = 10). The sample size indicates the number of embryos investigated. Sample size were not predetermined using statistical analyses. No blinding was performed for any analysis.

---

**Figure S1.**
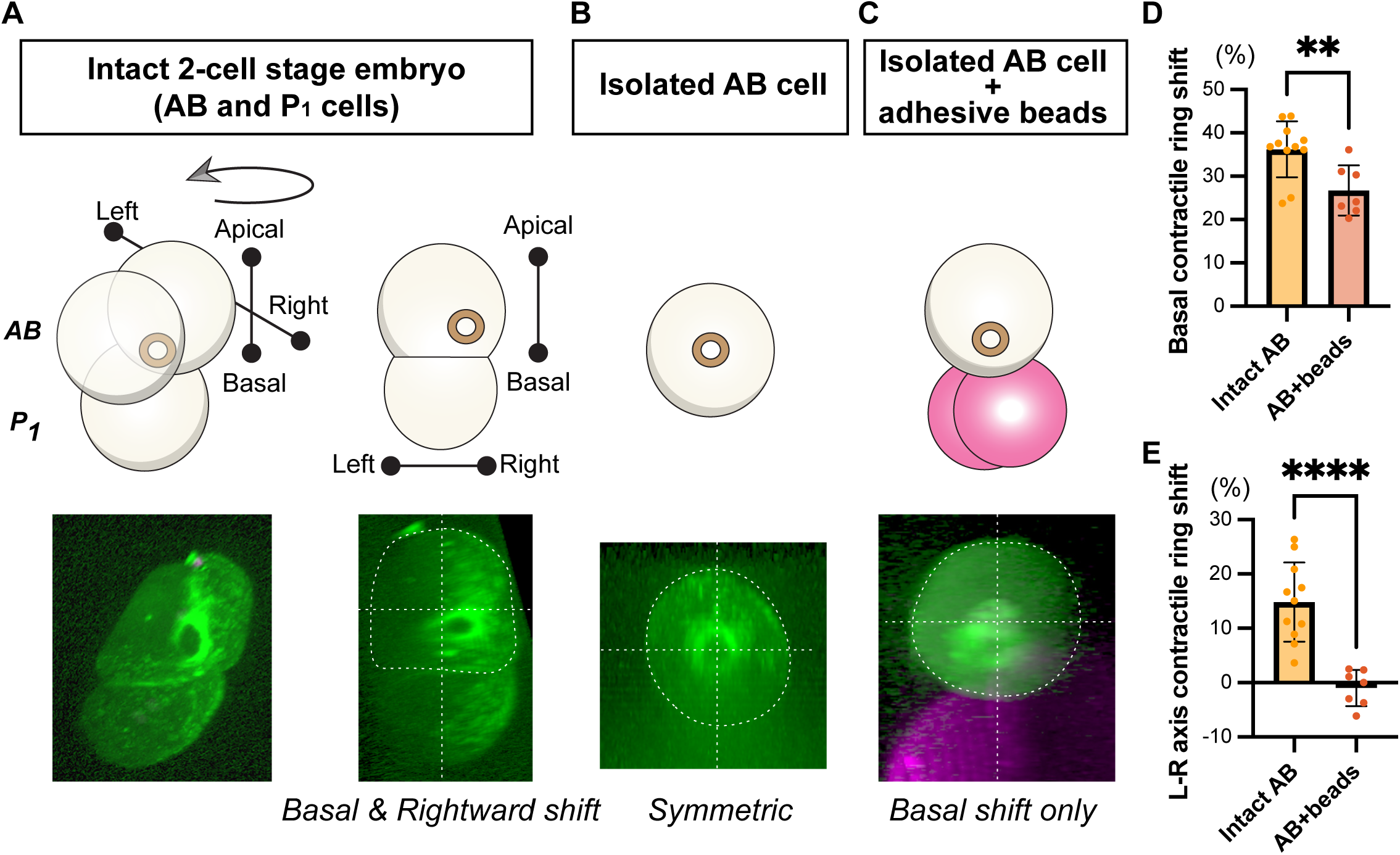
L-R asymmetric cytokinesis during 2nd mitosis of *C. elegans* embryo. (**A**) L-R Asymmetric contractile ring closure during cytokinesis of the AB blastomere at the 2-cell stage. The contractile ring position is shifted toward the basal (cell-cell contact) and right side from the center (bottom right panel). (**B**) As shown in Hsu et al., (2023), the contractile ring exhibits symmetric closure when separated from the neighboring P1 blastomere. (**C**) As shown in Hsu et al., (2023), isolated AB blastomere restores asymmetric contractile ring closure toward the basal side when attached by the adhesive beads. These beads were coated with positively charged Rhodamine, enabling electrostatic interaction with the negatively charged plasma membrane, generating adhesive forces. After re-analyzing our published dataset, we found that the contractile ring lost the L-R asymmetry. (**D-E**) Contractile ring shift along the L-R axis (D) and toward the basal side (E). Basal shift is observed in both intact and bead-attached AB cell. On the other hand, the shift along the L-R axis is diminished in the bead-attached AB blastomere. Green and magenta are NMY-2::GFP and adhesive beads, respectively. Data are obtained by re-analysis of dataset collected in Hsu et al., (2023). All the figures are newly prepared and not duplicated from the original work. Error bars: 95% confidence intervals. P-values: Welch’s t-test. “****” and “**” indicate p < 0.0001 and p < 0.01, respec-tively.

**Figure S2.**
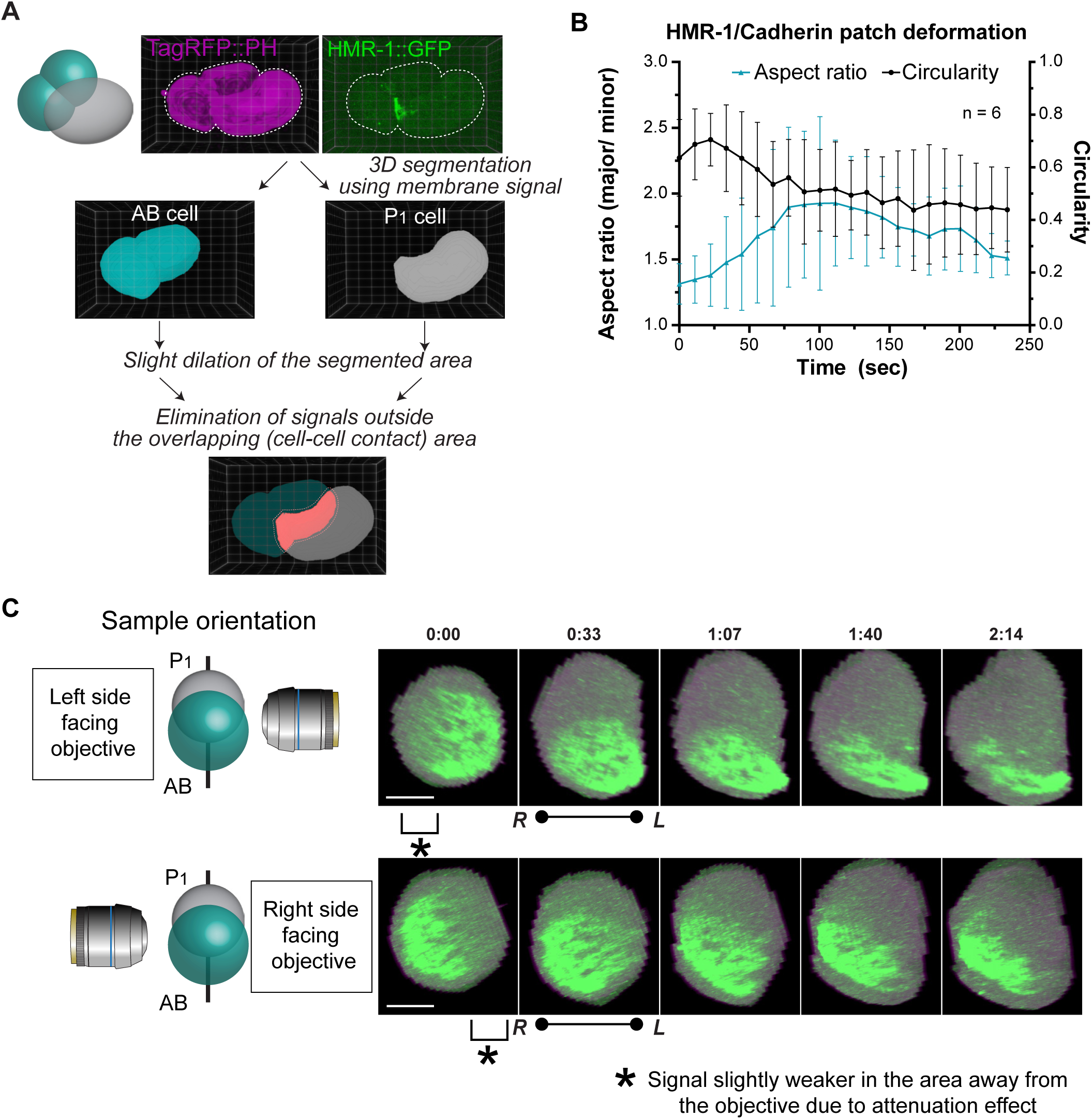
Viaualization and quantification of HMR-1::GFP at the cell-cell contact area. (**A**) Image analysis pipeline for isolating the contact area HMR-1::GFP signal. Embryos expressing TagRFP::PH (plasma membrane) and HMR-1::GFP (cadherin) were used. AB and P1 cell regions were segmented using morphological segmentation algorithm. The obtained segmented regions were slightly dilated, and signals outside the overlapping area were eliminated by filling zero values. (**B**) Quantification of the HMR-1 patch deformation process. The circularity and aspect ratio of the HMR-1 patch, viewed from the anterior region, are plotted relative to time after cytokinesis onset. (**C**) Effects of sample orientation on signal intensity along the imaging axis. Embryos were imaged compression-free and randomly oriented. Due to the attenuation effect, slices further from the objective lens exhibit lower signal intensity (asterisks). However, in both orientations, the same direction of HMR-1 patch twisting was observed. Times are relative to cytokinesis onset. Error bars indicate 95% confidence intervals.

**Figure S3.**
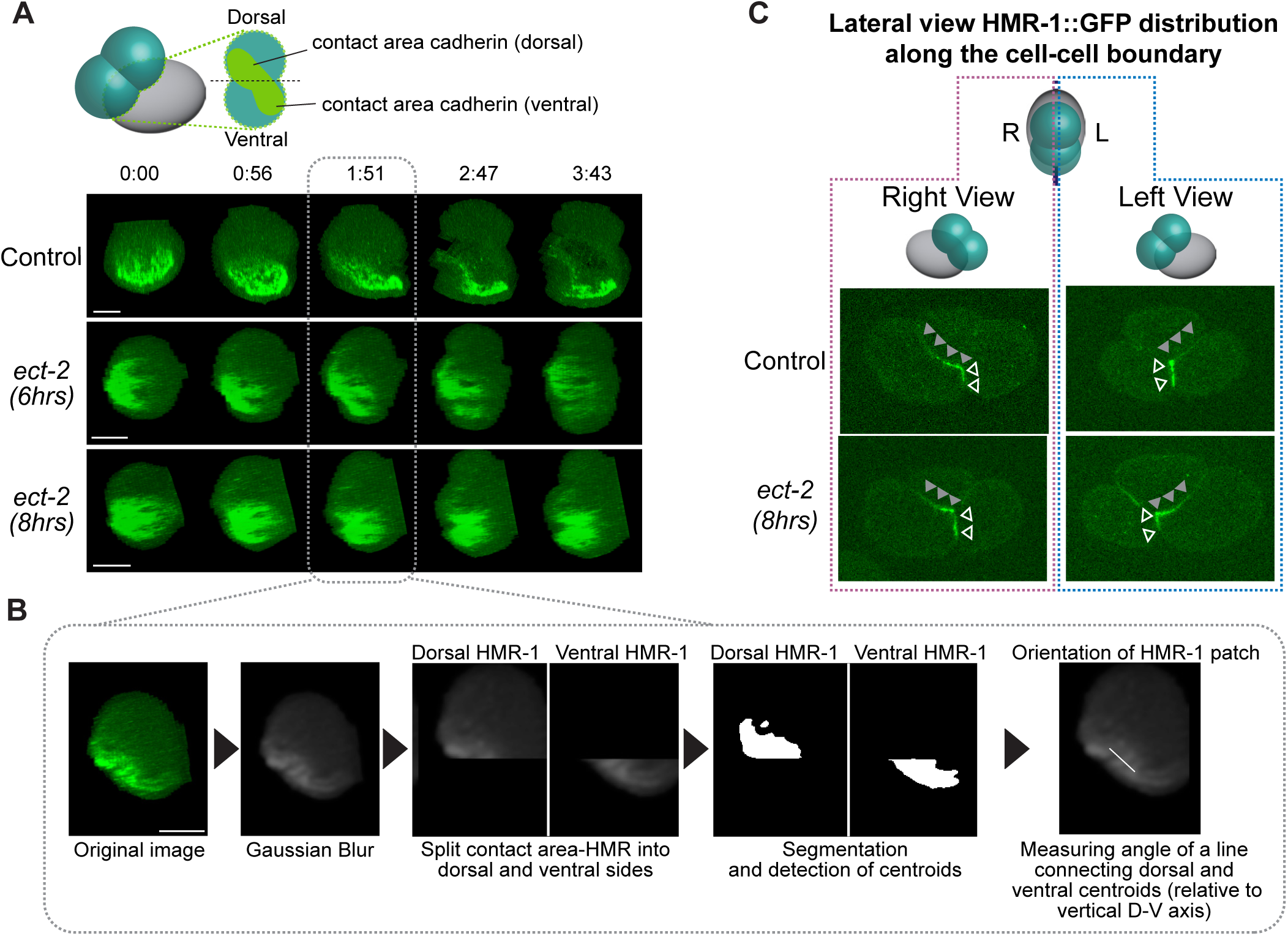
Effects of *ect-2(RNAi)* on HMR-1 patch twisting and how they were measured. (**A**-**B**) HMR-1::GFP signal at the AB-P1 contact area over time. Time is relative to cytokinesis onset (A). The quantification method for HMR-1 patch orientation (B). The HMR-1 signal was divided into dorsal and ventral parts based on the position of the cleavage furrow, as illustrated in the schematic in (A) and the images in (B). The dorsal and ventral HMR-1 patches were segmented, and their centroids were measured. The orientation of a line passing through the dorsal and ventral centroids were measured relative to the vertical, D-V axis. See also the schematic in Figure 2D. (**C**) HMR-1::GFP signal observed from the lateral sides of embryos. Images for the control are identical to those in Figure 1B for comparison. In *ect-2(RNAi;8hrs)* embryos, the difference in signal intensities between the dorsal (gray arrowheads) and ventral (white arrowheads) regions was lost.

**Figure S4.**
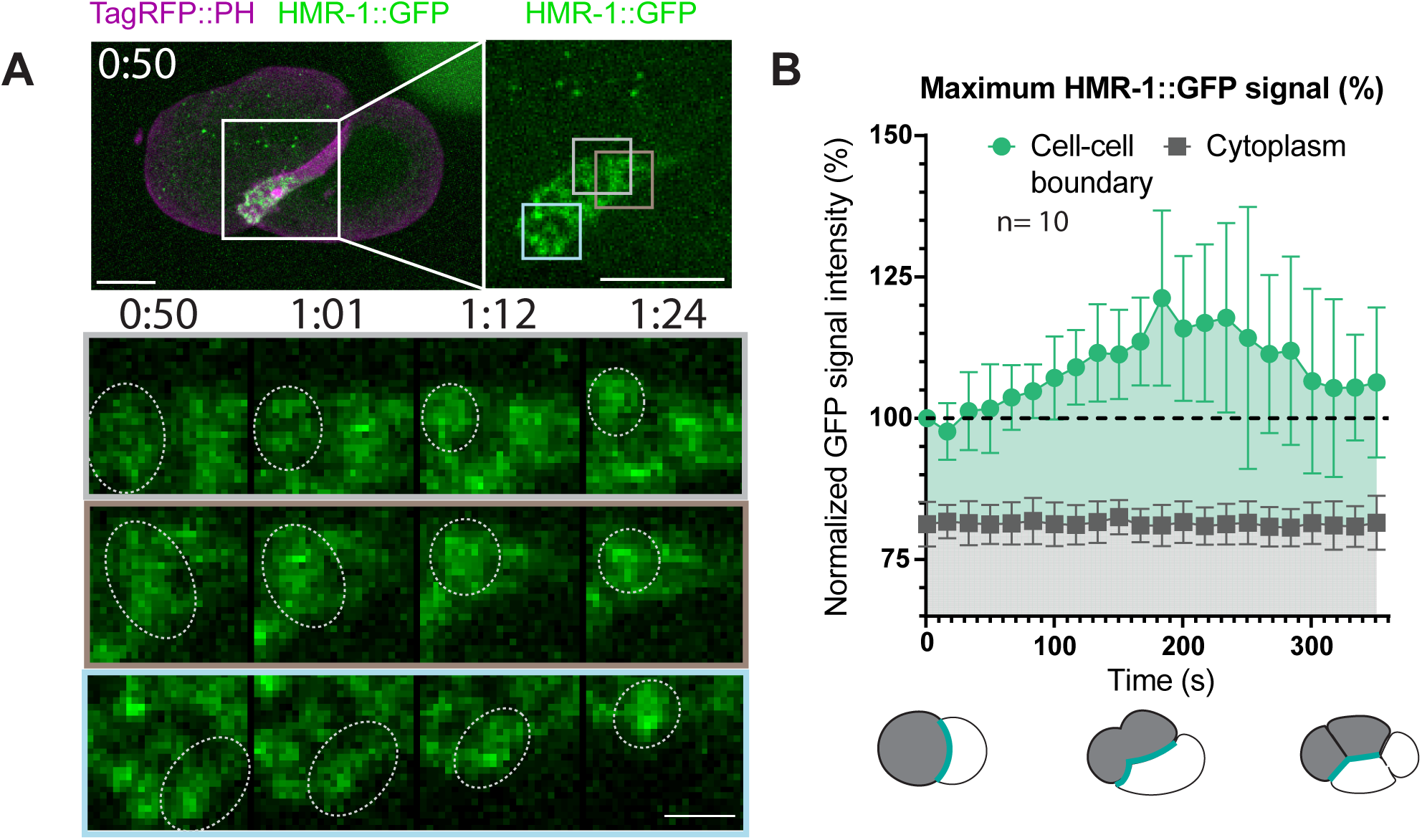
Coalescence of HMR-1 foci during their movement at the cell-cell contact membrane. (**A**) Movement of HMR-1::GFP foci at the AB-P1 contact area. Three independent coalescence events are highlighted in the indicated regions (colored boxes). (**B**) Maximum signal intensity of HMR-1::GFP during cytokinesis. Maximum intensity was measured using lateral view images, and both cytoplasmic and cell-cell boundary signals were quantified. The cell-cell boundary signal intensity at cytokinesis onset was set to 100%. Time is relative to cytokinesis onset. Error bars indicate 95% confidence intervals.

